# Optimized RNA-targeting CRISPR/Cas13d technology outperforms shRNA in identifying essential circRNAs

**DOI:** 10.1101/2020.03.23.002238

**Authors:** Yang Zhang, Tuan M. Nguyen, Xiao-Ou Zhang, Tin Phan, John G. Clohessy, Pier Paolo Pandolfi

## Abstract

Circular RNAs (circRNAs) are widely expressed, but their functions remain largely unknown. To study circRNAs in a high-throughput manner, short hairpin RNA (shRNA) screens^1^ have recently been used to deplete circRNAs by targeting their unique back-splicing junction (BSJ) sites. Here, we report frequent discrepancies between shRNA-mediated circRNA knockdown efficiency and the corresponding biological effect, raising pressing concerns about the robustness of shRNA screening for functional circRNAs. To address this issue, we leveraged the CRISPR/Cas13d system^2^ for circRNAs functional screenings. We optimized a strategy for designing single guide RNAs to deplete circRNAs. We then performed shRNA and CRISPR/Cas13d parallel screenings and demonstrated that shRNA-mediated circRNAs screening yielded a high rate of false positives phenotypes, while optimized CRISPR/Cas13d led to the identification of *bona-fide* functional circRNAs. Collectively, we developed a specific and reliable approach to functionalize circRNAs in a high-throughput manner.

## Main

Circular RNAs (circRNAs) are covalently closed, single stranded transcripts, which are produced by back-splicing of precursor mRNAs (pre-mRNAs). Thousands of circRNAs have been discovered across species with cell-type- and tissue-specific expression patterns ^3-6^. However, the functional repertoire of circRNAs remains mostly uncharacterized to date, which is mainly due to the unique properties of circRNAs and limitations of current approaches in circRNA studies^7, 8^. The rapid development of CRISPR-Cas9 based genomics screens has dramatically enhanced speed and precision of functional characterization of both coding genes and linear non-coning RNAs^9^. Nevertheless, the majority of circRNAs are generated from protein-coding genes^10^, and hence the sequences of circRNAs are completely overlapping with their cognate linear RNAs processed from the same pre-mRNA. Such features of circRNAs largely limit the application of Cas9 and its variants-mediated gene manipulations in understanding the functional relevance of circRNAs. Knockout of circRNAs is another loss of function (LOF) assay to study the function of circRNAs. And it could be achieved by depleting the complementary sequences (CSs) in flanking introns^11^, as the biogenesis of circRNAs are enhanced by RNA pairing of intronic CSs^12, 13^. However, the complexity of complementary sequence-mediated exon circularization^13, 14^ makes it difficult to apply this approach to annotate the functions of circRNAs at genome-wide scale. Therefore, although it is known that RNA interference (RNAi) has widespread non-specific transcripts silencing^15, 16^, RNAi-mediated degradation is still the major modality to date to silence circRNAs by targeting the unique BSJ site of circRNAs. Unfortunately, the requirement of designing shRNA/siRNA targeting BSJ sites limits the possibility to utilize multiple shRNAs/siRNAs with distinct coverage to rule out the potential off-target effects^7^. Recently, shRNA-based functional screen has been employed to understand circRNA essentiality^1^. However, in our study, we observed frequent discrepancies between shRNA-mediated circRNA knockdown efficiency and the corresponding biological effect on cell proliferation (see below), raising concerns about the robustness of using RNAi to study the function of circRNAs. Thus, the development of additional methods to achieve specific and efficient knockdown of circRNAs remains an important priority. In this study, we developed a strategy of designing CRISPR/Cas13d gRNA to specifically and effectively silence circRNAs, which are a lot more complex to target than linear transcripts. We also leveraged the optimized system for high-throughput circRNA functional screenings and compared its precision with shRNA-based screenings. Our platform proved to be more robust than shRNAs in identifying *bona-fide* functional circRNAs.

Given the tissue-specific expression pattern of circRNAs, in this study, we focused on human hepatocellular carcinoma (HCC) related circRNAs. We re-analyzed total RNA sequencing (rRNA depleted RNA-seq) data of paired primary tumors and adjacent normal tissues from 20 HCC patients ^17^ with CIRCexplorer2 to determine circRNA expression (**Supplementary Fig. 1a,b, see details in Methods**). We found 134 highly expressed circRNAs, and among which, 20 differentially expressed circRNAs were conserved between human and mouse (**Supplementary Fig. 1b,c**). Top 10 conserved circRNAs (**Supplementary Fig. 1c**) were selected for further characterization. RT-PCR with divergent primers across the BSJ sites followed by Sanger sequencing confirmed the junction sites, and RNase R resistance confirmed the circular structure of 9 out of 10 circRNAs, except for circARHGAP5 (or circArhgap5 in mouse) (**Supplementary Fig. 1d-h**). Moreover, most circRNAs predominantly localized in the cytoplasm, except for two circRNAs (circFBXW4 and circUBE3A) that showed half nuclear distribution (**Supplementary Fig. 1i**).

To investigate the functions of these validated circRNAs, we performed cell proliferation assay upon knockdown of each circRNA with two sets of shRNAs (**Supplementary Fig. 2a**). Knockdown efficiency of each shRNA was confirmed by qRT-PCR (**Supplementary Fig. 2b**). Interestingly, we found that knockdown of two circRNAs, circASPH and circZNF292, led to significant decreased proliferation rate compared to control cells (**Supplementary Fig. 2c,d**). However, we also noticed a dramatic difference in growth between two individual shRNAs, despite comparable knockdown efficiency (**Supplementary Fig. 2c-e**). The inconsistency between shRNA knockdown efficiency and inhibition of cell proliferation compelled us to include additional experimental strategies to assess the potential essentiality of these circRNAs. Antisense LNA GapmerRs, which is considered to be more specific than siRNA^18, 19^, were used to target circASPH (**Supplementary Fig. 2f**). Surprisingly, knockdown of circASPH by LNAs led to no obvious difference in the proliferation rate compared to control cells (**Supplementary Fig. 2g**), raising the concern about the reliability of shRNA to assess circRNA essentiality.

To address the shRNA issue and develop a more reliable knockdown tool to study the function of circRNAs, we sought to leverage the CRISPR/Cas13d system for depleting circRNAs. CRISPR/Cas13d system is a recently developed RNA-guided, RNA-targeting CRISPR system, which has been used to mediate efficient and specific knockdown of diverse linear transcripts^2^. Two distinct guide RNA architectures^2^, pre-gRNAs and gRNAs were employed to target circRNAs (**Supplementary Fig. 3a**). We found that pre-gRNA mediated a more potent knockdown (**Supplementary Fig. 3a**). Compared to gRNA with fixed 22 nt spacer, the transcribed pre-gRNA is processed into ∼ 52 nt mature gRNAs, with a 30 nt 5’ direct repeat followed by a variable 3’ spacer that ranged from 14-26 nt in length^2^. Therefore, we speculated that different spacer lengths may confer different levels of circRNA knockdown. To test our prediction, we generated a series of constructs that expressed progressively shorter gRNAs with spacers ranging from 30 nt to 21 nt in length (**Fig. 1a**). We found that gRNAs having 24 nt to 30 nt of target complementarity showed comparable knockdown efficacy, whereas Cas13d effector (CasRx) cleavage activity decreased when paired with gRNAs containing spacer sequence shorter than 23 nt (**Fig. 1a,b, Supplementary Fig. 3b**). We also observed that gRNAs having more than 30 nt of target complementarity showed less efficient knockdown of circRNAs (**Fig. 1c, Supplementary Fig. 3c**). To further finalize the optimal spacer length, we decided to evaluate the specificity of gRNAs with either 24 nt spacer (hereafter referred to as 24 nt gRNA) or 30 nt spacer (hereafter referred to as 30 nt gRNA) by assessing their sensitivity to Watson-Crick mismatches at the gRNA-DNA interface. We generated a series of variants of 24 nt gRNAs and 30 nt gRNAs targeting circZKSCAN1 (**Fig. 1d,e**). These variants contained single mismatches or consecutive double mismatches at indicated positions. We found that efficient knockdown of circRNAs was, in general, less compatible to mismatches inserted in the middle region (**Fig. 1d,e**). Intriguingly, 24 nt gRNAs-mediated knockdown was more sensitive to both single and double mismatches compared to 30 nt gRNAs (**Fig. 1d,e**). These findings were also observed in another set of gRNAs targeting circZNF292 (**Supplementary Fig. 3d,e**), confirming the choice for a 24 nt spacer. Taken together, we successfully developed a strategy to generate CRISPR/Cas13d gRNAs with optimal efficacy and specificity for circRNA knockdown, and gRNAs with the 24 nt spacer design were used for subsequent experiments.

**Fig. 1.**
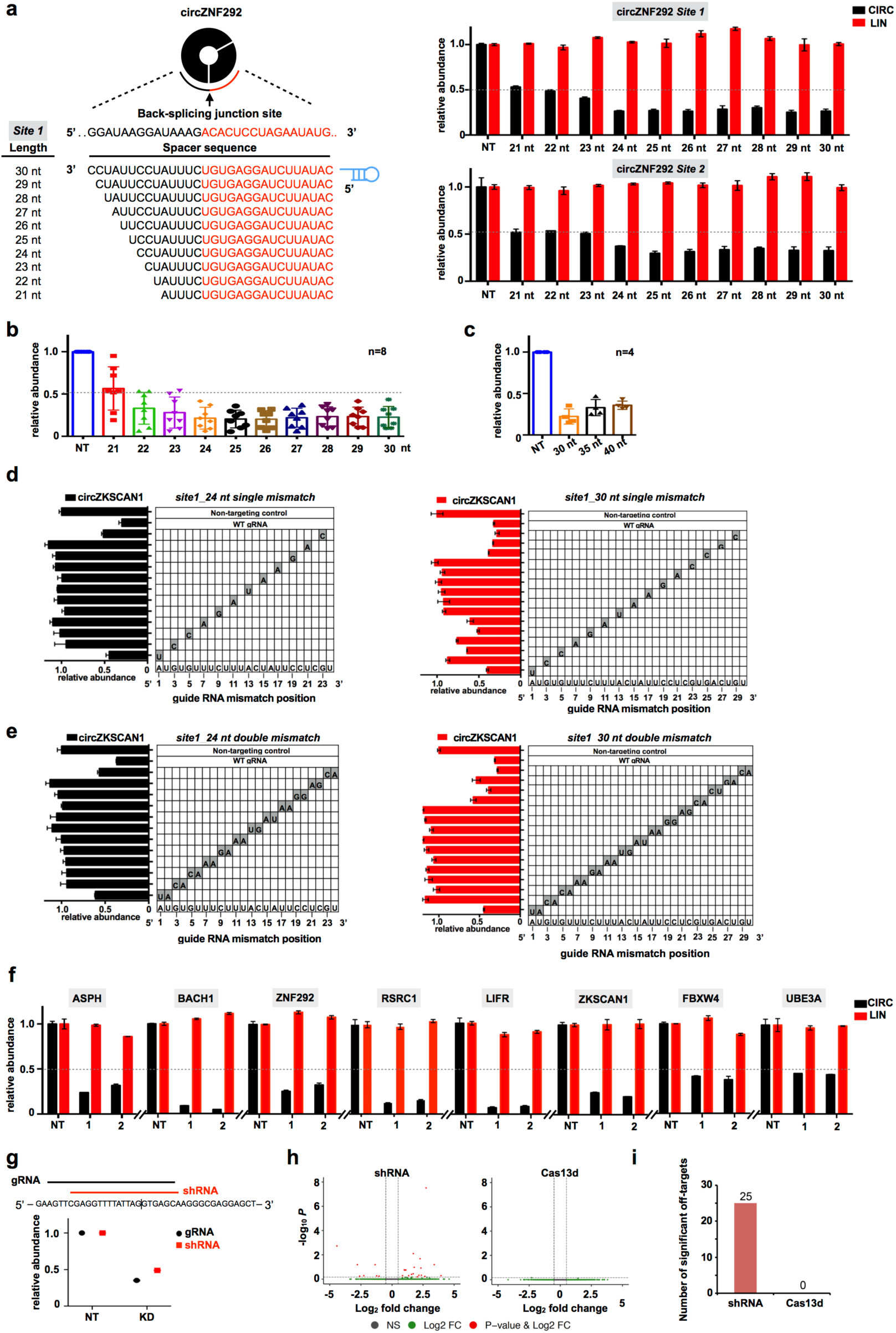
Optimization of CRISPR-Cas13d for circRNAs knockdown. **(a)** Schematic view of the length optimization of gRNAs targeting BSJ sites of circRNAs (left panel) and the corresponding knockdown efficiency with different lengths of gRNAs (right panel). Bar plots showing the relative expression of circZNF292 (CIRC) and its parental linear transcript (LIN) upon knockdown of circZNF292 with two gRNAs containing spacers ranging from 21 nt to 30 nt in length. NT, non-targeting. (**b)** Bar graph showing the cumulative knockdown efficiency of different length of gRNAs across multiple circRNAs (n=8 independent experiments, the data for each experiment can be found in Fig. 1a and Supplementary Fig. 3). NT, non-targeting. (**c)** Bar graph showing cumulative knockdown efficiency of longer gRNAs with 30 nt, 35 nt and 40 nt length spacers (n=4 independent experiments, the data for each experiment can be found in Supplementary Fig. 3). NT, non-targeting. (**d)** Knockdown of circZKSCAN1 evaluated with gRNAs containing 24 nt length spacer (left) or 30 nt length spacer (right) with single mismatch at varying positions across the spacer sequence. The gray boxes in the grids show the position of Watson-Crick transversion mismatches. The wild-type sequence is shown at the bottom of each grid. (**e)** Knockdown of circZKSCAN1 evaluated with gRNAs containing 24 nt length spacer (left) or 30 nt length spacer (right) with consecutive double mismatch at varying positions across the spacer sequence. The gray boxes in the grids show the position of Watson-Crick transversion mismatches. The wild-type sequence is shown at the bottom of each grid. (**f)** Optimized Cas13d system targeting 8 circRNAs, each with 2 gRNAs. qRT-PCR for circular and linear transcripts after knockdown of circRNAs in Huh7 cells. (**g)** Top, schematic drawing of circEGFP-targeting guide RNA sequence and spacer position-matched shRNA. Bottom, relative circEGFP knockdown by individual position-matched gRNA and shRNA. NT, non-targeting. (**h)** Volcano plots of differential transcript levels between circEGFP targeting and non-targeting shRNAs (left) or circEGFP-targeting CasRx and non-targeting guide (right) as determined by RNA sequencing. (**i)** Summary of significant off-target transcript perturbations by matched Cas13d gRNA and shRNA. The data shown are from one of two biological replicates with similar results, and error bars indicating the mean ± s.d. of three technical replicates.

To test our hypothesis concerning shRNA off-target effects for circRNA functional characterization, we applied the optimized CRISPR/Cas13d system to silence circASPH, which showed inconsistent growth phenotypes with different shRNAs as well as in comparison with the LNA knockdown method (**Supplementary Fig. 2c,f,g**). Huh7 cells with stably expressed CasRx were transduced with gRNAs containing 24 nt spacer sequences targeting BSJ site of circASPH. QRT-PCR and northern blot (NB) confirmed Cas13d-mediated circASPH knockdown without affecting the corresponding linear mRNA (**Supplementary Fig. 2h,i**). Similar to LNA-mediated circASPH knockdown, no obvious difference in the proliferation rate was observed between circASPH-silencing cells and control cells (**Supplementary Fig. 2j**), demonstrating the off-target effects of shRNA for circRNA knockdown, and further proving the reliability of Cas13d in assessing the function of circRNAs.

To further evaluate the range of efficiency of optimized Cas13d knockdown, we designed gRNAs targeting the same endogenous circRNAs that have been successfully silenced by shRNAs (**Supplementary Fig. 2b**). As revealed by qRT-PCR, gRNAs showed comparable knockdown activity to shRNA-mediated circRNAs degradation (**Fig. 1f**).

Importantly, we sought to optimize the targeting of circRNAs with nuclear distribution (circFBXW4 and circUBE3A). To this end, we employed and adapted a nuclear-localized version of CasRx (CasRx with nuclear localization signal, CasRx-NLS) ^2^. Indeed, gRNAs paired with CasRx-NLS mediated higher levels of knockdown than shRNAs (**Supplementary Fig. 3f, 2b**), suggesting that Cas13d is a more versatile tool to target circRNAs with different localization.

Next, to comprehensively assess and compare the specificity of Cas13d and shRNA for circRNA knockdown, position-matched gRNA and shRNA were used to target circEGFP (**Supplementary Fig. 3g**). Both methods achieved comparable levels of circEGFP knockdown (**Fig. 1g**). Since circEGFP is not endogenous to the cell, cells with circEGFP knockdown should have similar transcriptomic profiles to cells transduced with non-targeting shRNA or gRNA. We observed that compared to Cas13d, shRNAs showed higher variability between targeting and non-targeting conditions (**Fig. 1h**). Differential expression analysis indicated 25 significant off-targets in shRNA condition but none in Cas13d condition (**Fig. 1i**). Collectively, compared to shRNA, our optimized Cas13d showed comparable circRNA knockdown efficiency, but with high specificity.

We further evaluated the capacity of CRISPR/Cas13d system to screen for essential circRNAs in a high-throughput manner. We performed in parallel both Cas13d and shRNA screens in Huh7 cells in order to systematically compare the two systems’ abilities to identify circRNAs that are essential for cell growth (**see details in Methods**). Briefly, position-matched gRNAs and shRNAs were designed to target the BSJ sites of 134 highly expressed circRNAs (**Fig. 2a, Supplementary Fig. 4a**). gRNA and shRNA libraries were lentivirally infected into cells, and screened for gene essentiality over a 14-day period (**Fig. 2b**). PCR-amplified barcode-gRNAs or shRNAs from genomic DNA of cells before and after screening were subjected to deep sequencing. Overall, the read distribution of duplicated screens within each condition showed a high level of correlation for both gRNA and shRNA (**Fig. 2c, Supplementary Fig. 4b**). To identify the top hits from the screens, we processed our sequencing data using MAGeCK algorithm (v0.5.8). Gene set enrichment analysis (GSEA) showed that both gRNAs and shRNAs targeting positive controls (10 known essential linear transcripts^20^) were significantly enriched in the ranked list of negative selected gRNAs or shRNAs (**Fig. 2d**), suggesting that these two parallel screens performed as intended. However, compared to non-targeting control gRNAs in Cas13d library, non-targeting control shRNAs had a much higher level of variation (**Fig. 2e,f**). The observed high level of correlation between duplicated screens rules out the possibility that the high level of variation was due to technical variation. Therefore, this variation in non-targeting controls is more likely due to shRNA’s off-target effects. In contrast, non-targeting control gRNAs have a much narrower range of variation (**Fig. 2e,f**), confirming Cas13d’s high level of specificity. For circRNAs, MAGeCK identified 10 negatively selected circRNAs with statistical significance (false discovery rate (FDR)<0.25) from shRNA-based screen (**Fig. 2g**), including circASPH that was tested in Supplementary Fig. 2. Six of the remaining candidates were resistant to RNase R treatment (**Supplementary Fig. 5a**), confirming their existence as circRNAs. These 6 circRNA candidates were further validated for their essentiality in conferring growth in Huh7 cells. We performed cell proliferation assays upon knockdown of each circRNA with five individual shRNAs presented in the library. Knockdown efficiency of each shRNA was confirmed by qRT-PCR, and most of them resulting in >60% reduction of circRNA abundance (**Fig. 3a,e, Supplementary Fig. 5b,f,j,n**). However, we also noticed that several circRNA-targeting shRNAs decreased the counterpart linear transcripts as well, especially for circNPEPPS (**Supplementary Fig. 5b**). Notably, similar to circASPH (**Supplementary Fig. 2c**), the inconsistency between shRNA-mediated circRNA knockdown efficiency and effect on cellular proliferation rate was detected in 6 out of 6 testing circRNAs (**Fig. 3a-h, Supplementary Fig. 5b-q**), suggesting a wide-spread off-target effects of shRNA in circRNA knockdown. To confirm that the shRNA screen identified false positive essential circRNAs, we used position-matched gRNAs from the Cas13d screening library to target the same circRNAs. Consistent with our prediction, no obvious change in the cell proliferation rate was observed in Cas13d mediated circRNA knockdown cells compared to control cells, despite the comparable level of knockdown of the target circRNAs to shRNAs (**Fig. 3a-h, Supplementary Fig. 5b-q**). Taken together, these data indicate the high false positive rate of shRNA screens for identifying essential circRNAs.

**Fig. 2.**
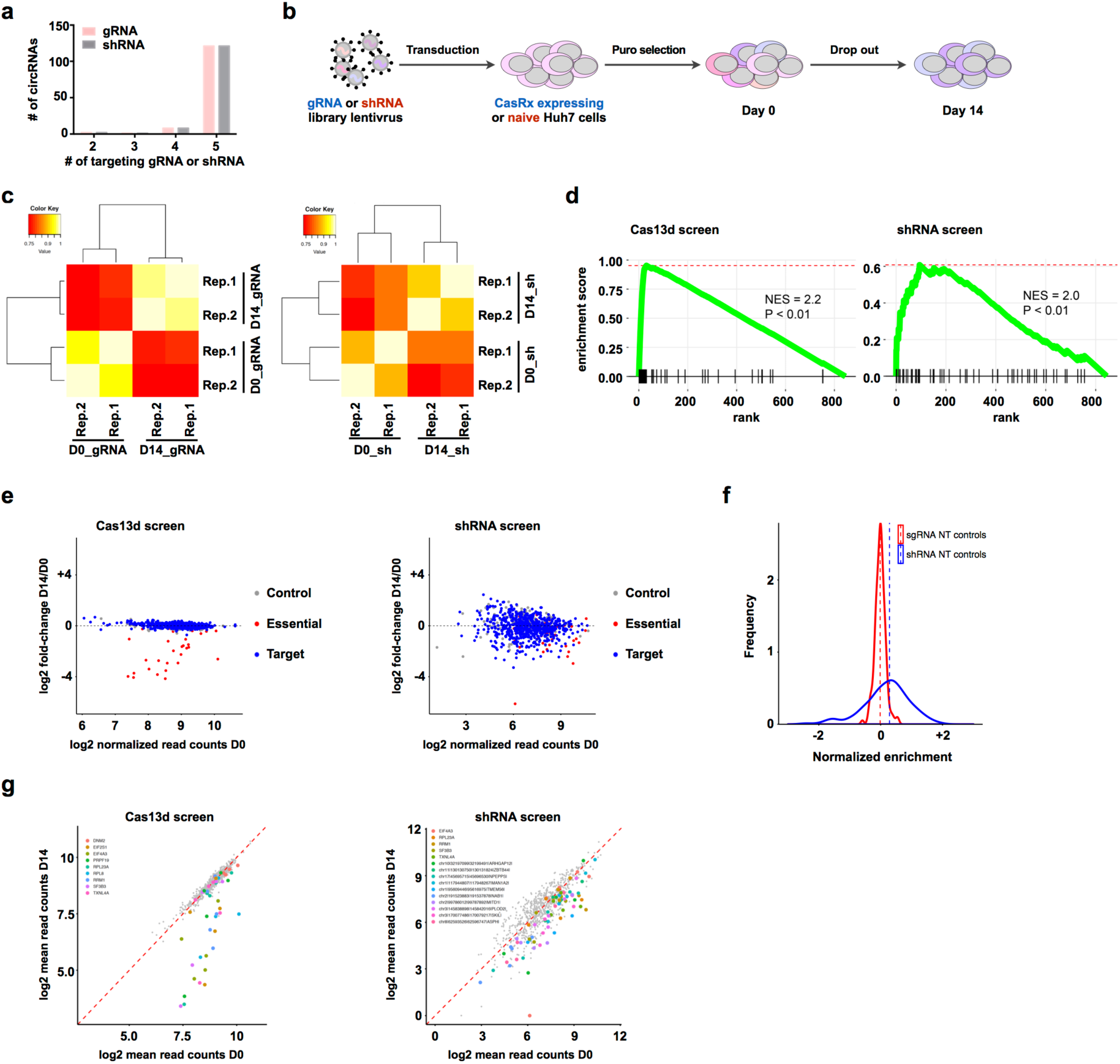
Systematic comparison of CRISPR-Cas13d and shRNA functional screens for circRNAs. **(a)** Number of gRNAs and shRNAs per circRNA in the library. **(b)** Schematic view of screens. Cas13d and shRNA lentivirus libraries were infected into CasRx stably expressed Huh7 cells or naive Huh7 cells separately and selected by puromycin treatment (time zero). Puromycin-resistant cells were further cultured for 14 days. Genomic DNA was extracted at indicated time points and library representation was determined by deep-sequencing. **(c)** Correlation heatmap showing the Pearson correlation coefficient between the levels of gRNAs/shRNAs in biological replicates of time zero samples (D0) and 14-day enrichment samples (D14) for Cas13d screen (left) and shRNA screen (right). **(d)** Gene set enrichment analysis (GSEA) revealed essential genes are enriched in negative selections for Cas13d screen (left) and shRNA screen (right). Essential genes serve as positive controls. The degree of enrichment is measured as normalized enrichment score (NES). **(e)** Scatterplots showing log_2_-transformed fold-change of gRNA/shRNA normalized read counts in D14 vs. D0 for Cas13d screen (left) and shRNA screen (right). Control, non-targeting controls; Essential, positive controls targeting known essential genes; Target, circRNAs highly expressed in HCC. **(f)** Histograms representing the relative distribution of non-targeting control gRNAs and shRNAs. **(g)** Scatterplots showing negatively selected gRNA/shRNAs and corresponding genes from Cas13d screen (left) and shRNA screen (right) with FDR < 0.25. CircRNAs are indicated with genomic locations and the host gene name at the end (e.g. chr10|32197099|32199491|ARHGAP12|). Positive controls only have gene names without genomic location (e.g. EIF4A3)

**Fig. 3.**
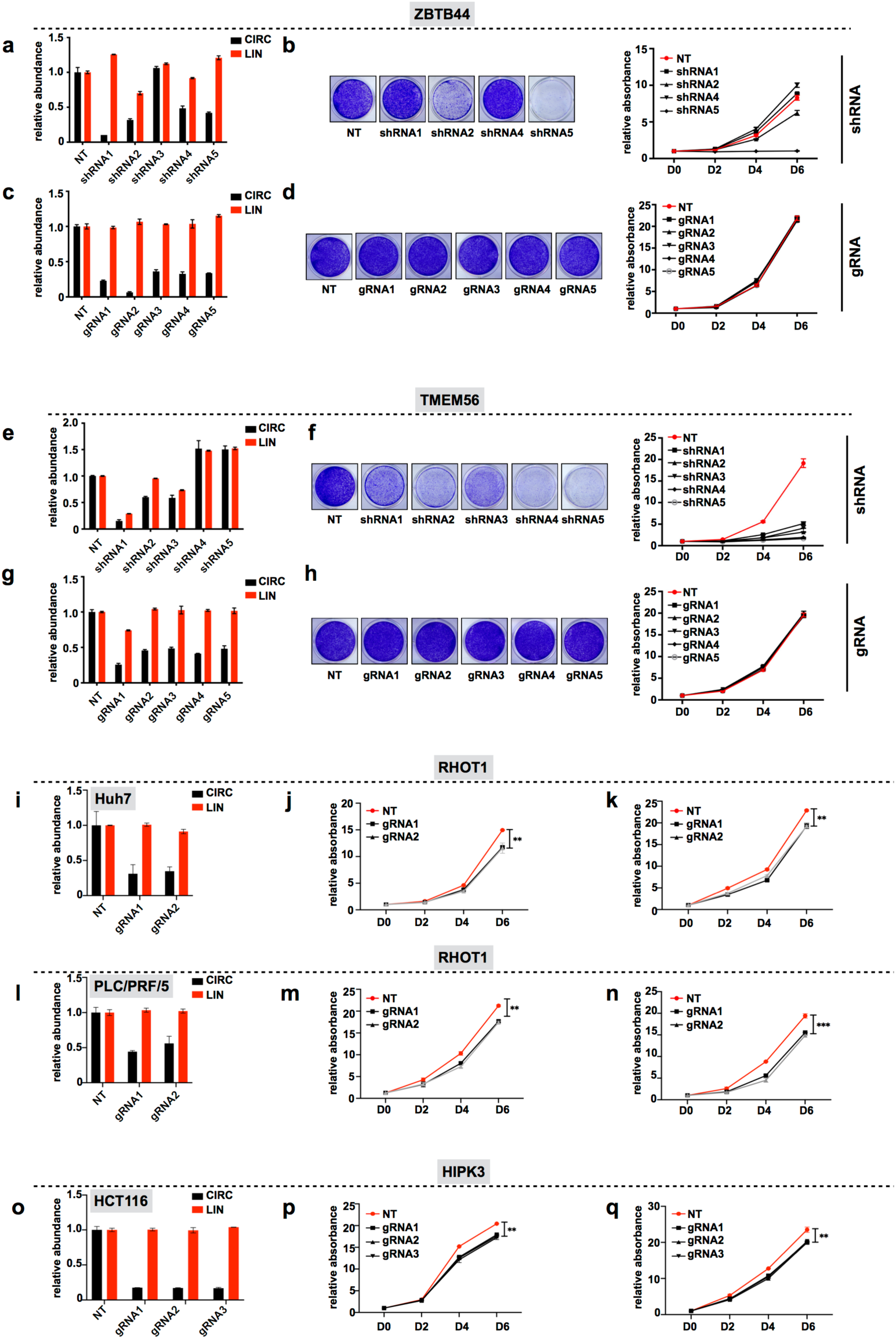
Cas13d identified *bona-fide* essential circRNAs, whereas shRNA screen identified false-positive targets. **(a,c)** Relative expression levels of circZBTB44 and its parental mRNA upon knock-down of circZBTB44 by shRNAs (a) and gRNAs (c) in human Huh7 cells. **(b, d)** Proliferation rates of control and shRNA-mediated (b) or Cas13d-mediated (d) circZBTB44-silenced Huh7 cells. The number of cells was detected upon staining with crystal violet, and representative pictures are shown on the left, while the proliferation curves are shown on the right. **(e, g)** Relative expression levels of circTMEM56 and its parental mRNA upon knock-down of circTMEM56 by shRNAs (e) and gRNAs (g) in human Huh7 cells. **(f, h)** Proliferation rates of control and shRNA-mediated (f) or Cas13d-mediated (h) circTMEM56-silenced Huh7 cells. The number of cells was detected upon staining with crystal violet, and representative pictures are shown on the left, while the proliferation curves are shown on the right. **(i, l)** Relative expression levels of circRHOT1 and its parental mRNA upon knock-down of circRHOT1 by gRNAs in Huh7 cells (i) or PLC/PRF/5 cells (l). **(j, m)** Proliferation rates of control and circRHOT1-silenced Huh7 cells (j) or PLC/PRF/5 cells (m). The number of cells was detected upon staining with crystal violet. **(k, n)** Proliferation rates of control and circRHOT1-silenced Huh7 cells (k) or PLC/PRF/5 cells (n) assessed using a CCK-8 kit at indicated days. **(o)** Relative expression levels of circHIPK3 and its parental mRNA upon knock-down of circHIPK3 by gRNAs in HCT116 cells. **(p)** Proliferation rates of control and circHIPK3-silenced HCT116 cells. The number of cells was detected upon staining with crystal violet. **(q)** Proliferation rates of control and circHIPK3-silenced HCT116 cells assessed using a CCK-8 kit at indicated days. The data shown are from one of two biological replicates with similar results, and error bars indicating the mean ± s.d. of three technical replicates. \**p* < 0.05, \*\**p* < 0.01, \*\*\**p* < 0.001 (unpaired student’s *t* test). ns, not significant.

We analyzed Cas13d screening sequencing data using the same algorithm. Positive control gRNAs were significantly enriched as expected given the essential roles of their targets, whereas none of gRNAs targeting circRNAs dropped out during the screen (**Fig. 2g**). One possibility could be the small size of the library (targeting 134 circRNAs), limiting the probability to identify essential circRNAs. However, none of the false positive candidates identified in the shRNA screen was detected in the Cas13d screen, suggesting that our Cas13d screening platform has a much lower false positive rate compared with the conventional shRNA screening platform.

To evaluate the ability of Cas13d to identify and credential *bona-fide* essential circRNAs, we next sought to use our optimized Cas13d system to target previously reported essential circRNAs. For instance, circRHOT1 knockout by depleting flanking CSs suppressed HCC’s cell proliferation^21^. To determine if Cas13d can validate the essentiality of circRHOT1 in HCC cells, we silenced its expression in two HCC cell lines, Huh7 and PLC/PRF/5 cells (**Fig. 3i,l**). Functional experiments using crystal violet staining and CCK8 assay demonstrated that circRHOT1 knockdown inhibited cell proliferation (**Fig. 3j,k,m,n**), which further proved the ability of Cas13d to identify functional circRNAs. circHIPK3 is another circRNAs that is known to promote proliferation of the colon cancer cell line HCT116 ^22^. circHIPK3 was silenced by Cas13d in HCT116 cells (**Fig. 3o**), and three individual gRNA-mediated knockdown confirmed the effect of circHIPK3 in cell proliferation (**Fig. 3p,q**), demonstrating the capability of Cas13d to assess the essentiality of circRNAs. Collectively, our optimized Cas13d platform is effective in identifying *bona-fide* essential circRNAs with a much smaller false positive rate compared with shRNAs.

Different approaches have been developed to study the function of linear transcripts, whereas functional analysis on circRNAs has been challenging. The development of an appropriate tool to knockdown circRNAs without affecting their cognate linear RNAs is key to understanding the functional and biological relevance of circRNAs. In this study, we developed a more reliable method to elucidate the function of circRNAs and strategy for designing gRNAs for the Cas13d system in order to achieve specific and efficient knockdown of circRNAs. In general, Cas13d paired with gRNAs containing 24 nt spacers did silence on-target circRNAs with high efficiencies and showed reduced silencing effects at closely matched off-target sites. Compared to the widely used RNAi approach, optimized Cas13d knockdown of circRNAs was comparable to RNAi knockdown efficiency, but with substantially reduced off-target effects, making it well-suited for systemic evaluation of circRNA functions. From side-by-side comparison of CRISPR/Cas13d and shRNA screens, we found that the abilities of the two libraries to detect known essential linear genes were similar, but for circRNAs, shRNA screen yielded a much high rate of false positive phenotypes. Moreover, our results demonstrate that optimized Cas13d can validate the phenotypes of previously reported *bona-fide* functional circRNAs, demonstrating the capability and reliability of Cas13d in unraveling the relevance of truly functional circRNAs. Furthermore, our study is the first to develop a high-throughput screening system with CRISPR-Cas13d for manipulating circRNAs, and paves the way for future studies using the system for screening functional noncoding RNAs that are more problematic to screen with the DNA-editing CRISPR-Cas9 system.

Our results raise in turn several important questions to be addressed in future experiments. Although we have proven the ability of Cas13d to identify essential circRNAs, we did not obtain any circRNA positive hits from the Cas13d screening. There are three possible explanations for this result. The first possibility is that we designed gRNAs to target 134 highly expressed circRNAs, which only cover a minor fraction of expressed circRNAs in Huh7 cells. As a consequence, we may have missed truly functional circRNAs. The second possibility is that many circRNAs may function cooperatively instead of individual molecules. CircRNAs have been implicated in the regulation of innate immunity^23, 24^, and a recent study revealed that many cytoplasmic circRNAs create a molecular reservoir of PKR to prevent aberrant activation of innate immunity in uninfected cells^25^. Therefore, knocking down individual circRNAs may not lead to significant phenotypes. The third possibility is that circRNAs may have functions other than regulation of cell proliferation as studied in this work such as responding to external stimuli or stresses. Further Cas13d-based functional screens will be required to improve our understanding of the functional relevance of circRNA.

In summary, we have developed and optimized a novel CRISPR/Cas13d based approach that dramatically reduces off-target background noise in both the screening and validation of truly functional circRNAs. This novel approach will tremendously facilitate the annotation of the functional circRNA landscape in both physiological processes and disease pathogenesis.

## Acknowledgement

We thank all of the members of the Pandolfi lab for their support and critical discussions. This work was supported by an R35 NCI (CA197529-01) grant to P.P.P.

## Author Contributions

Y.Z. and P.P.P. conceived of and designed experiments. Y.Z., T.M.N., and T.P. performed experiments. X.-O.Z. performed the bioinformatics analysis of HCC patients seq data and designed the libraries. T.M.N. performed computational analysis of screens and circEGFP-seq analysis. Y.Z., T.M.N., X,-O.Z. drafted the original manuscript. Y.Z., T.M.N., J.G.C., and P.P.P. reviewed and edited the paper. P.P.P. supervised the project.

## Methods

### Computational Pipeline for HCC circRNAs annotation

We used the CIRCexplorer2 pipeline^14^ to annotate expressed circRNAs in HCC patients as candidate circRNA targets. In brief, sequencing reads of ribo-depleted total-RNA-seq datasets^17^ (GSE77509) of 40 samples from 20 HCC patients (each with one pair of primary tumor and adjacent normal tissue sample) were aligned to the GRCH37/hg19 human reference genome by STAR (parameters: -- chimSegmentMin 10) to identify chimeric junction reads. Chimeric junction reads were then filtered and compared against the UCSC gene annotation (updated at 2016/9/17) to quantify the expression of circRNAs using CIRCexplorer2, and 134 expressed circRNAs were selected with RPM (reads per million mapped reads) ≥ 0.1 in all the 20 HCC patients as candidate circRNAs for further screening. For further experimental validation, a subset of circRNAs were selected according to the following criteria: a) average fold change ≥ 1.5 between primary tumors and adjacent normal tissues from 20 HCC patients, b) conserved between human and mouse (at least two unique reads in mouse liver samples). This filtering yielded 20 circRNAs, and top 5 up-regulated and top 5 down-regulated circRNAs were then selected for experimental validation.

### Cell Culture and treatment

Human cell lines including Huh7, HepG2, Hep3B, SK-Hep1, SNU475, SNU423, SNU387, PLC/PRF/5, HCT116 and HEK293T cells were purchased from American Type Culture Collection (ATCC). Huh7, SK-Hep1, PLC/PRF/5, HCT116 and HEK293 cells were maintained in DMEM supplemented with 10% FBS at 37 °C with 5% CO_2_. HepG2 and Hep3B cells were maintained in MEM supplemented with 10% FBS at 37 °C with 5% CO_2_. SNU475, SNU423 and SNU387 cells were maintained in RPMI 1640 supplemented with 10% FBS at 37°C with 5% CO_2_. To generate Huh7, PLC/PRF/5 and HCT116 cells with stable expression of CasRx, the cells were transduced by EF1a-CasRx (no NLS-RfxCas13d)-2A-EGFP (modified from Addgene #109049) lentivirus, and CasRx positive cells were then collected through cell sorting for EGFP marker. Antisense LNA GapmerRs were synthesized at QIAGEN and were transfected into Huh7 cells with Lipofectamine 2000 (Invitrogen) according to the manufacturer’s instructions with a concentration of 50 nM.

### Plasmids Construction

The lentiviral gRNA and pre-gRNA expressing backbones were constructed by cloning the human U6 promoter and CasRx gRNA or pre-gRNA scaffold (Addgene #109053, #109054) into lentiGuide-Puro (Addgene, #52963) by replacing its original U6-gRNA cassette. To construct individual gRNA or pre-gRNA expressing vector, the annealing pairs of oligonucleotides (Invitrogen) harboring complementary sticky ends were ligated to BsmBI-cleaved gRNA or pre-gRNA backbones. The oligonucleotides for shRNA were cloned into the pLKO.1-TRC vector (Addgene #10878) using AgeI/EcoRI. To construct circEGFP expressing plasmid, a partial EGFP sequence was inserted into lentiviral backbone with two complementary sequences in the flanking intron. All the plasmids used in this study will be available from the corresponding author upon reasonable request.

### RNA Isolation, qRT-PCR, RT-PCR and Northern Blotting

Total RNA from cultured cells with different treatments was extracted with Trizol Reagent (Invitrogen) according to the manufacturer’s protocol. For qRT-PCR and RT-PCR, the cDNA synthesis was carried out using SuperScript IV (Invitrogen) with random hexamers. QPCR was done using SybrGreen reaction mix (Applied Biosystems) and StepOnePlus™ real-time PCR system (Applied Biosystems). The relative expression of different sets of genes was normalized to GAPDH mRNA level. Northern blotting was carried out according to the manufacturer’s protocol (DIG Northern Starter Kit, Roche). RNA was loaded on denatured PAGE gels. Digoxigenin (Dig) labeled antisense probes were generated using T7 RNA polymerase by *in vitro* transcription with the RiboMAX Large Scale RNA Production System (Promega).

### RNase R treatment

To enrich circRNA isoforms, 10 µg total RNA was diluted in 20 µl of water with 4U RNase R/µg and 2 µl enzyme buffer (Epicentre), then incubated at 37 °C for 3 h. 10 µg total RNA incubated with buffer only was used as controls. Both RNase R treated and untreated RNAs were further subjected to Trizol extraction and followed by qRT-PCR or RT-PCR.

### Nuclear/Cytoplasmic RNA Fractionation

Cellular fractionation in Huh7 cells was performed as previously described^26^. Briefly, 2 × 10^7^ Huh7 cells were used for nuclear/cytoplasmic RNA fractionation. Cell pellet was suspended by gentle pipetting in 200 µl lysis buffer (10 mM Tris ph8.0, 140 mM NaCl, 1.5 mM MgCl2, 0.5% Igepal, 40U/ml Recombinant RNasin Ribonuclease Inhibitor), and incubated on ice for 10 min. During the incubation, one tenth of the lysate was added to 1 ml Trizol for total RNA extraction. The rest of the lysate was centrifuged at the 1000 rpm for 3 min at 4 °C to pellet the nuclei, and the supernatant was the cytoplasmic fraction. Fractionated RNAs from the same amount of cells were used for cDNA synthesis and qRT-PCR.

### Cell Proliferation assay

Cell proliferation was measured using crystal violet staining or CCK-8 kit. For crystal violet staining, cells were seeded at a concentration of 4 × 10^4^ cells (Huh7 and PLC/PRF/5 cells) or 3 × 10^4^ cells (HCT116 cells) per well in a 12-well plate and cultured for 6 days in complete medium (DMEM plus 10% FBS) at 37 °C. Cells were fixed with 10% formalin at indicated days and stained with 0.1% crystal violet. Crystal violet was then solubilized with 10% acetic acid, and their absorbance was measured using SpectraMax iD3 Multi-Mode Microplate Readers. For CCK-8 assay (Abcam, ab228554), 3x 10^3^ cells were seeded in 96-well plates and cultured for 6 days in complete medium (DMEM plus 10% FBS) at 37 °C. At indicated time points, cells were incubated with 10 µl of CCK-8 assay solution in each well for 2 h at 37 °C. The absorbance values at 460 nm were then measured using SpectraMax iD3 Multi-Mode Microplate Readers.

### Lentivirus preparation and transduction

Low passage HEK293T cells were transfected with Lipofectamine 2000 (Thermo Fisher Scientific) and Cas13d plasmid or guide RNA expressing plasmid plus pMDG.2 and psPAX2 packaging plasmids. After 24 h, the medium was changed to prewarmed DMEM medium. Viral supernatant was harvested 48 h later, and cellular debris was filtered out using Millipore’s 0.45 µm PVDF filter. To assess the knockdown ability of individual gRNA or shRNA, Huh7 cells stably expresseing Cas13d or naive Huh7 cells were infected with gRNA or shRNA lentivirus. After 24 h post-transduction, the medium was changed to fresh medium with 2 µg/ml puromycin. After 5 days post-transduction, total RNAs were harvested for further analysis.

### RNA sequencing and analysis

For specificity analysis, RNA sequencing was performed on rRNA-depleted total RNA from cells with Cas13d and shRNA-mediated circEGFP knockdown. Total RNA was extracted from cells infected with lentiviruses carrying knockdown constructs using Trizol. rRNA depleted total RNA-Seq libraries were prepared by the Molecular Biology Core Facilities (MBCF) at Dana-Farber Cancer institute (DFCI). RNA-Seq libraries were sequenced on an Illumina NextSeq instrument with at least 10M reads per library. RNA-Seq reads were aligned and quantified with Salmon (v0.13.1)^27^ using default parameters for paired-end reads with --validateMappings flag. Human reference transcriptome available in Ensemble portal (ftp://ftp.ensembl.org/pub/release-95/fasta/homo_sapiens/cdna/) were indexed for Salmon alignment and quantification. Transcript per million (TPM) values, averaged from biological replicates, were transformed to log scale for expression correlation. To find differentially expressed genes, raw transcript counts generated with Salmon were imported into DESeq2 (v1.26.0)^28^ for count normalization and differential expression analysis. Genes with no read count in at least 1 sample were not included in the analysis. Only genes that had a log2 differential expression greater than 0.5 or less than −0.5 and a false discovery rate < 0.68 were reported to be significantly differentially expressed.

### shRNA and gRNA library design

To perform functional screening, 134 highly expressed circRNAs in HCC were selected. For each circRNA candidate, all the possible 21 nt shRNA target sequences were extracted from the back-splice junction sequence (40-nt long with 20 nucleotides at each side of back-spliced exons), and scored by siDirect version 2.0 (http://sidirect2.rnai.jp/design.cgi) and GPP web portal (https://portals.broadinstitute.org/gpp/public/). To remove possible off-target sequences, all shRNA candidates were aligned back to the human transcriptome (GENCODE V19) permitting 3 mismatches with bowtie (parameters: -n 3 −l 5 --norc -y -a). The shRNA sequences were selected based on the following criteria: high on-target sequence score, high coverage of the BSJ site, high complexity of the library. The final shRNA library contained 646 shRNAs targeting 132 circRNAs, and most circRNAs had 5 shRNAs (for two circRNAs, none of the shRNAs passed the filters). To generate a comparable gRNA library, gRNA sequences were designed by extending each shRNA from 21 nucleotides to 24 nucleotides and filtered by off-target blast. To evaluate the efficiency of our screens, cell-essential genes (CRISPR score < –1 and adjusted p-value < 0.05 in all examined cell lines) were downloaded from a CRISPR/Cas9-based genome-wide negative selection screening study^20^, and ten top cell-essential genes with the lowest mean CRISPR scores were selected as positive controls. For each positive control gene, five top shRNAs with highest adjusted score were downloaded from the Genetic Perturbation Platform (https://portals.broadinstitute.org/gpp/public/). To minimize off-target effects, all control shRNAs were aligned back to the human transcriptome (GENCODE V19) permitting 3 mismatches using bowtie (parameters: -n 3 -l 5 --norc -y -a). Corresponding gRNAs targeting positive essential genes were designed by extending shRNA sequences to 24 nt oligonucleotides. In addition, 150 random intergenic regions (RefSeq gene annotations updated at 2017/5/28) in the fly genome (dm6) were selected as negative controls.

### Construction of the Cas13d gRNA and shRNA libraries and libraries screening

Cas13d gRNA library were synthesized as 94-mer oligonucleotides (CustomArray), caccgaacccctaccaactggtcggggtttgaaacNNNNNNNNNNNNNNNNNNNNNNNNttttttaagcttggcgtaactagatcttgagacaa (N indicates the 24 nt spacer sequence), and amplified by PCR as a pool using the following primers: tatatatcttgtggaaaggacgaaacaccgaacccctaccaactggtcggggtttgaaac (Forward), cttttaaaattgtggatgaatactgccatttgtctcaagatctagttacgccaagc (Reverse). shRNA library were synthesized as 92-mer oligonucleotides (CustomArray), ggaaaggacgaaacaccggNNNNNNNNNNNNNNNNNNNNNctcgagNNNNNNNNNNNNNNNNNNNNNtttttgaattctcgacctcgagaca (N indicated 21 nt target sequence), and amplified by PCR as a pool using the following primers: taacttgaaagtatttcgatttcttggctttatatatcttgtggaaaggacgaaacaccgg (Forward), cccccttttcttttaaaattgtggatgaatactgccatttgtctcgaggtcgagaattc (Reverse). The PCR product was purified and then cloned into gRNA-expressing or shRNA-expressing vector using NEBuilder HiFi DNA Assembly Master Mix (NEB #E2621). 100 ng product was then transformed into Endura ElectroCompetent cells according to the manufacturer’s directions. Clones were scraped off the LB plates and plasmid DNA was extracted using PureLink™ HiPure Plasmid Maxiprep Kit (ThermoFisher, K210007). The libraries were submitted for next generation sequencing to confirm the coverage and diversity of gRNA and shRNA libraries. The lentivirus of gRNA or shRNA library was produced by co-transfection of library plasmids with two viral packaging plasmids psPAX and pMD2.G into HEK293 cells using Lipofactamine 2000 (Invitrogen). Huh7 cells were transduced with lentivirus libraries at multiplicity of infection (MOI) ∼ 0.3. Replicated transductions were performed. 24 h after transduction, cells were cultured with fresh medium containing 2 µg/ml puromycin. After two days of puromycin selection, genomic DNA was extracted as Day 0. In the screen, cells were passaged every 3 days, and maintained a coverage of >500 cells per gRNA or shRNA. After 14 days screening, genomic DNA was extracted for replicated samples. gRNA and shRNA inserts were amplified using 10 different NGS-lib-Forward primers paired with Reverse primers containing unique barcode. gRNA and shRNA distribution was determined by next-generation sequencing. The libraries were sequenced on the Illumina MiSeq according to the user manual (Harvard Medical School Biopolymers Facility, Boston).

### Computational analysis of screens

The screening sequencing data was analyzed using MAGeCK (v0.5.8)^29^. MAGeCK “count” command was used to generate read counts of all samples as previously described^30^. Briefly, raw read counts were normalized with DESeq2 then rlog transformed for generation of correlation heatmaps and PCA plots (**Fig. 3d** and **Supplementary Fig. 4b**). MAGeCK “test” command was used to identify the top negatively and positively selected circRNAs as previously described^30^. MAGeCK estimates the level of negative (or positive) selection of each circRNA by comparing the rankings of all gRNAs or shRNAs targeting that circRNA with a null model, where all gRNAs/shRNAs are distributed uniformly in the ranked list. The α-Robust Rank Aggregation (α-RRA) algorithm was used to calculate the “RRA score” of each circRNA to describe the degree of negative (or positive) selection. The *P* value of the RRA score was computed by permuting all circRNAs, and adjusted for multiple comparison correction with the Benjamini-Hochberg method. A detailed description of the algorithm is reported in the original study^29^.

### Gene set enrichment analysis

Preranked GSEA of gRNAs and shRNAs for positive controls (known essential genes) was conducted using the fgsea (v1.12.0) R package.

