## Supplemental Material for "Optimized RNA-targeting CRISPR/Cas13d technology outperforms shRNA in identifying essential circRNAs"

**Inventory of Supplemental information**

I. Supplemental Figures

Supplementary Fig. 1 is related to main Fig.1.

Supplementary Fig. 2 is related to main Fig.1.

Supplementary Fig. 3 is related to main Fig.1.

Supplementary Fig. 4 is related to main Fig.2.

Supplementary Fig. 5 is related to main Fig.3.

**Supplemental Figures and Figure Legends**

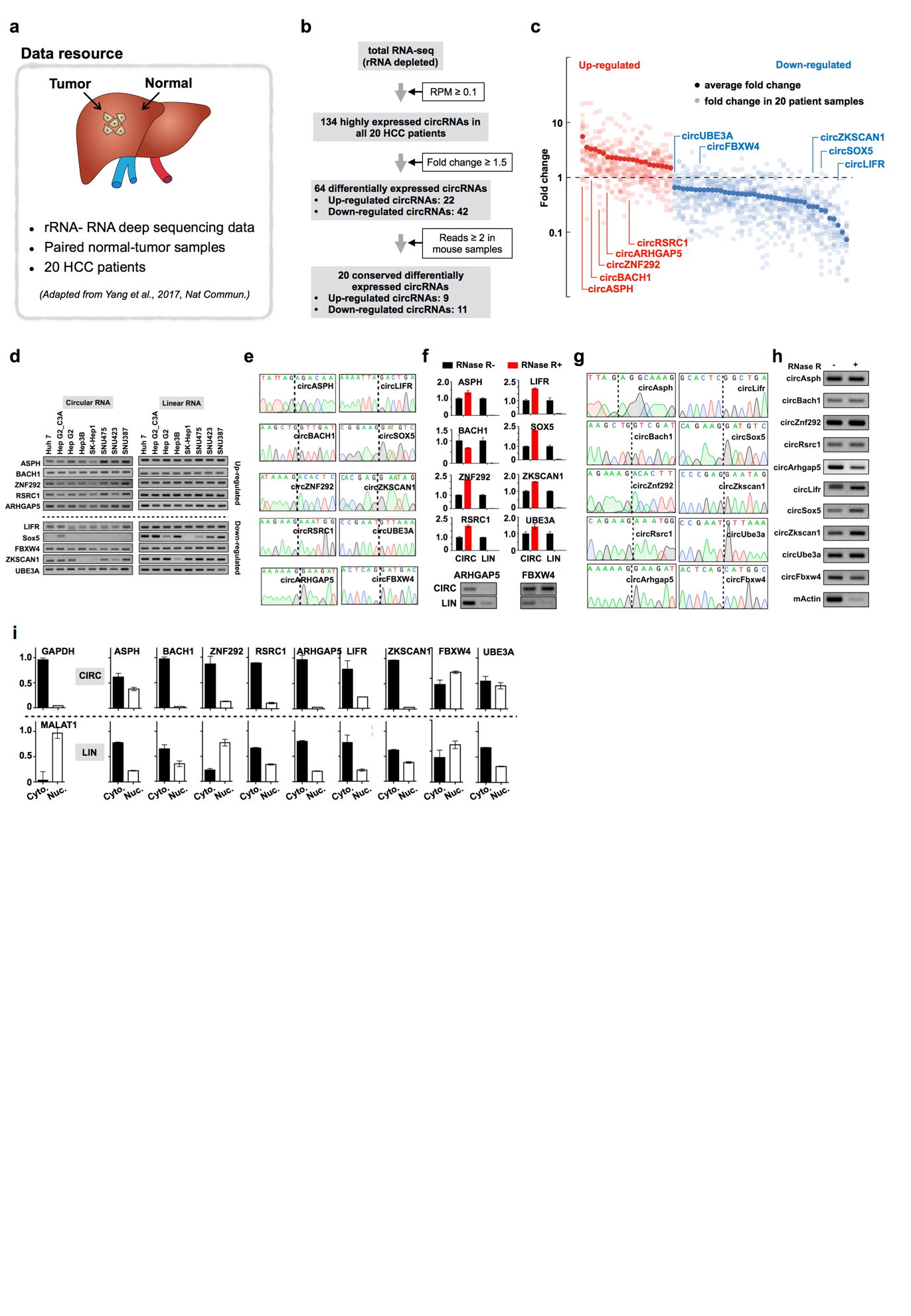

**Supplementary Fig. 1 | Identification and characterization of circRNAs in HCC. (a)**

Schematic drawing of data resource used for identification of circRNAs in HCC. **(b)** The schematic diagram shows the computational pipeline for the systematic identification of circRNAs in samples from HCC patients. **(c)** Scatterplots showing fold change of differentially expressed circRNAs in samples from HCC patients. X axis, 64 circRNAs were rank-ordered by differential expression between primary tumor samples and paired-adjacent normal tissues. Light colored dots represent the fold change of circRNAs in each paired samples from 20 HCC patients , and dark colored dots represents the average fold change. Y axis, fold change of the expression level of individual circRNAs. A subset of circRNAs validated herein are labeled. **(d-e)** Expression of human circRNAs were validated by RT-PCR (d), followed by Sanger sequencing (e). RT-PCR validation of 10 conserved differentially expressed circRNAs in 8 human HCC cell lines: circRNAs and their corresponding linear transcripts were amplified with divergent and convergent primers. Left panel, agarose electropherogram of circRNA PCR products. Right panel, agarose electropherogram of the corresponding linear mRNA PCR products (d). PCR products were subjected to Sanger sequencing, back-splicing junction sites are indicated by dash lines (e). **(f)** RNase R validation of 10 selected circRNAs. circRNAs together with the corresponding linear mRNAs were amplified by qRT-PCR or RT-PCR from cDNA prepared from RNA non-treated or treated with RNase R. **(g-h)** Validation of 10 conserved circRNAs in mouse liver samples. BSJ sites were indicated by dashed lines (g). circRNAs were amplified by RT-PCR from mouse liver cDNA prepared from non-treated or treated with RNase R. Mouse actin mRNA was used as a negative control (h). **(i)** Subcellular localization of human circRNAs. Bar plots represent relative abundance of RNAs in nuclear and cytoplasmic fractions. The relative distribution of GAPDH and MALAT1 transcripts, predominantly localized to the cytoplasm and nucleus, respectively, confirmed a successful cellular fractionation. Error bars in **f** ,**i** indicating the mean ± s.d. of three technical replicates.

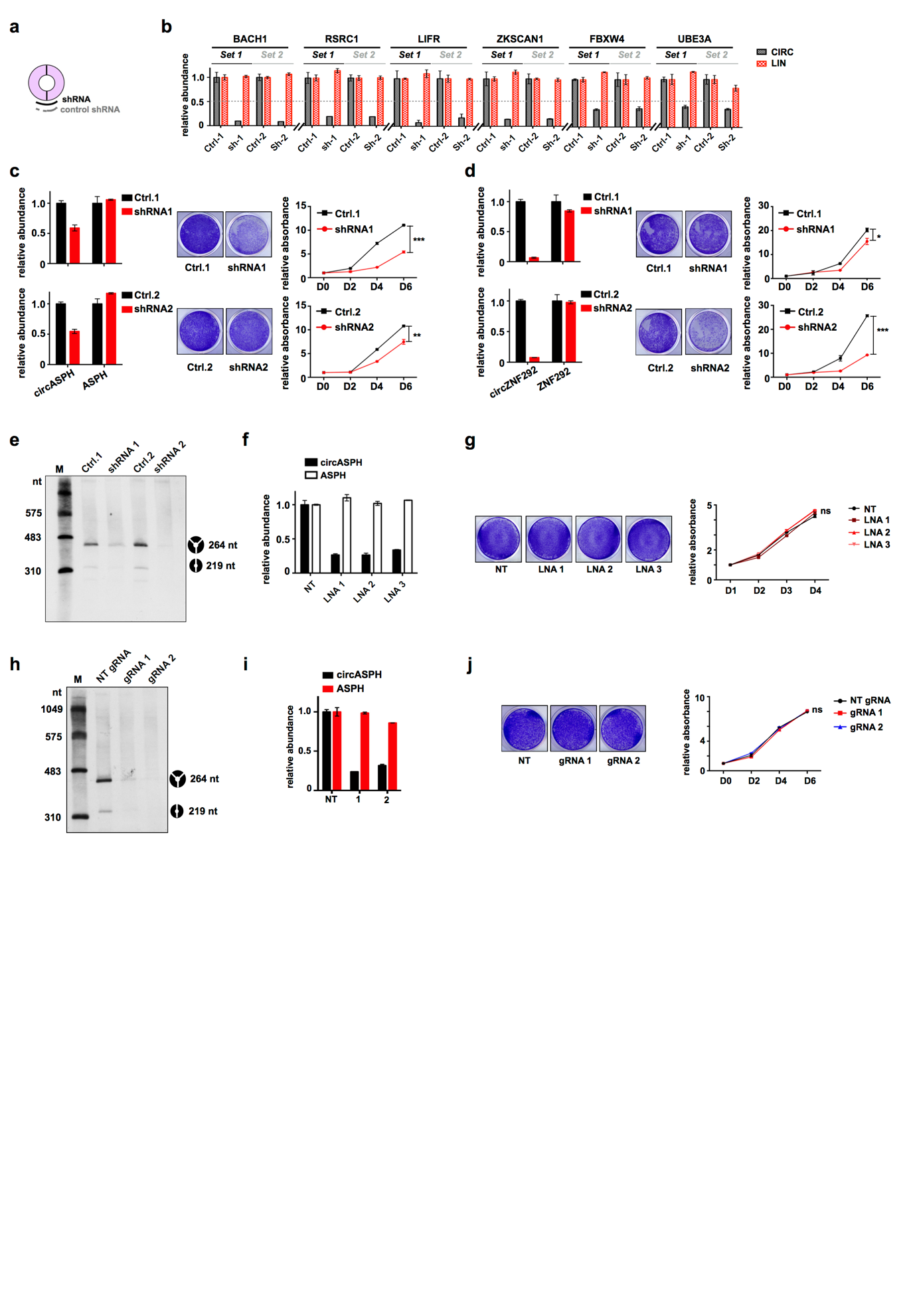

**Supplementary Fig. 2 | Targeting conserved HCC circRNAs with shRNAs. (a)** Schematic drawing of the strategy to silence the expression of circRNAs with shRNAs targeting the BSJ sites. ShRNAs with half-scrambled sequence were used as controls. **(b)** Relative expression levels of circRNAs and their parental genes upon knock-down of circRNAs by two sets of shRNAs in Huh7 cells. **(c)** Knockdown of circASPH in Huh7 cells. Relative expression levels of circASPH and its parental mRNA upon knock-down of circASPH by two sets of shRNAs in human Huh7 cells (left panel). Proliferation rates of control and circASPH-silenced Huh7 cells (right panel). **(d)** Knockdown of circZNF292 in Huh7 cells. Relative expression levels of circZNF292 and its parental mRNA upon knock-down of circZNF292 by two sets of shRNAs in human Huh7 cells (left panel). Proliferation rates of control and circZNF292-silenced Huh7 cells (right panel). **(e)** NB revealed the relative abundance of two isoforms of circASPH after knockdown circASPH with two shRNAs. circASPH contains an alternative internal exon, which leads to the production of two isoforms of circASPH shared the same BSJ site. **(f)** Relative expression levels of circASPH and its parental mRNA upon knock-down of circASPH by three LNAs in Huh7 cells. LNA, locked nucleic acid. **(g)** Proliferation rates of control and LNA-mediated circASPH-KD Huh7 cells. The number of cells was detected upon staining with crystal violet, and representative pictures are shown on the left, while the proliferation curves are shown on the right. **(h)** NB revealed the relative abundance of circASPH after knockdown circASPH with two gRNAs. **(i)** Relative expression levels of circASPH and its parental mRNA upon knock-down of circASPH by CasRx paired with gRNAs. **(j)** Proliferation rates of control and Cas13d-mediated circASPH-KD Huh7 cells. The number of cells was detected upon staining with crystal violet, and representative pictures are shown on the left, while the proliferation curves are shown on the right. The data shown are from one of two biological replicates with similar results, and error bars indicating the mean ± s.d. of three technical replicates. **p* < 0.05, ***p* < 0.01, ****p* < 0.001 (unpaired student’s *t* test). ns, not significant.

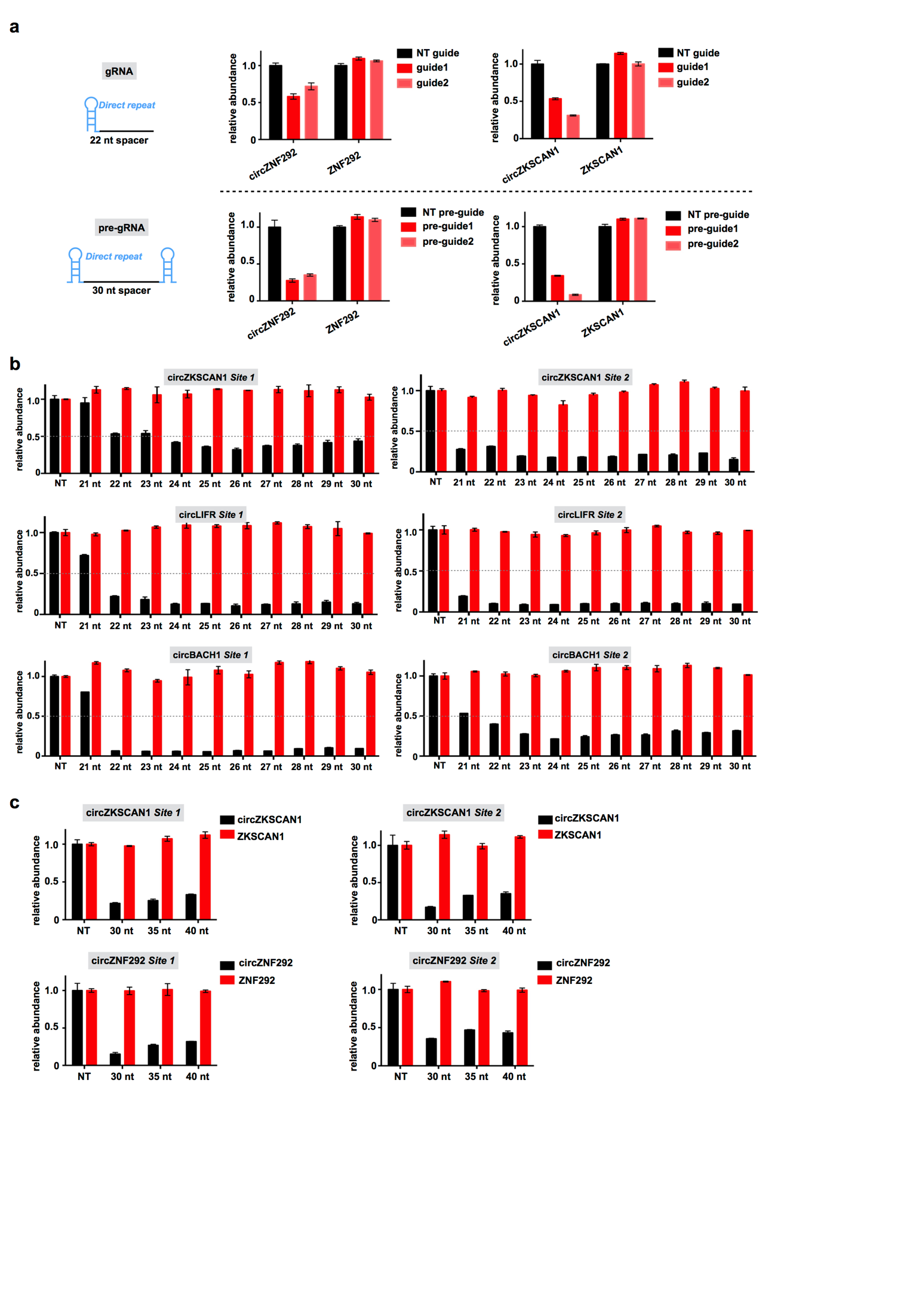

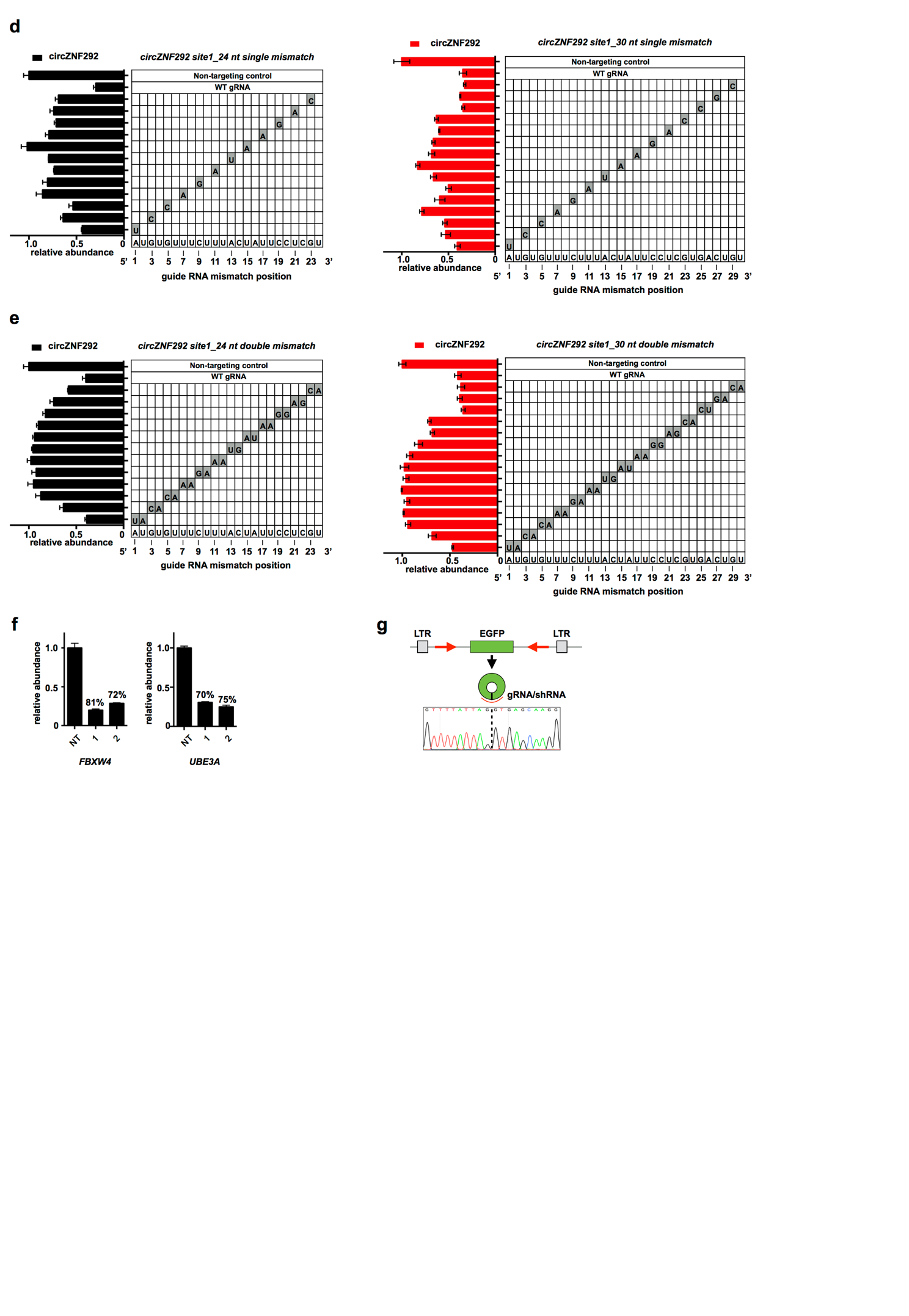

**Supplementary Fig. 3 | Adaptation of CRISPR-Cas13d system to silence circRNAs. (a)** Comparison of gRNA and pre-gRNA for knockdown of endogenous circRNAs reveals pre-gRNA to be the more effective guide RNA architecture. Schematic for constructs expressing guide RNAs. pre-gRNA, unprocessed guide RNA containing a single 30 nt spacer sequence flanked by 2 full-length 36 nt Direct repeats. gRNA, predicted mature guide RNA with a single 30 nt processed Direct repeat and 22 nt spacer sequence. NT, non-targeting. **(b)** Bar plots showing the relative expression of three more circRNAs and their linear parental RNAs upon knockdown of circRNAs with different length of gRNAs targeting two regions of BSJ site of each circRNA. NT, non-targeting. **(c)** Bar plots showing the relative expression of circZKSCAN1 and its parental linear mRNA (top) or circZNF292 and its parental linear mRNA (bottom) upon knockdown of circRNAs with gRNAs containing 30 nt, 35 nt and 40 nt length spacers. **(d)** Knockdown of circZNF292 evaluated with gRNAs containing 24 nt length spacer (left) or 30 nt length spacer (right) with single mismatch at varying positions across the spacer sequence. The gray boxes in the grids show the position of Watson-Crick transversion mismatches. The wild-type sequence is shown at the bottom of each grid. **(e)** Knockdown of circZNF292 evaluated with gRNAs containing 24 nt length spacer (left) or 30 nt length spacer (right) with consecutive double mismatch at varying positions across the spacer sequence. The gray boxes in the grids show the position of Watson-Crick transversion mismatches. The wild-type sequence is shown at the bottom of each grid. **(f)** qRT-PCR for relative expression of circFBXW4 (left) and circUBE3A (right) after knockdown circRNA with NLS-CasRx. Numbers above the bar represent the knockdown level. NLS-CasRx, CasRx with nuclear localization signal. **(g)** Schematic drawing of circular EGFP expression vector (Top). Partial sequence of EGFP was inserted into circRNA expression vector with fully complementary sequences in the flanking introns to facilitate the biogenesis of circEGFP. The complementary sequences are indicated with red arrow to show the polarity. Bottom, Sanger sequencing result confirms the BSJ site of circEGFP, as indicated by black dash line. The data shown are from one of two biological replicates with similar results, and error bars indicating the mean ± s.d. of three technical replicates.

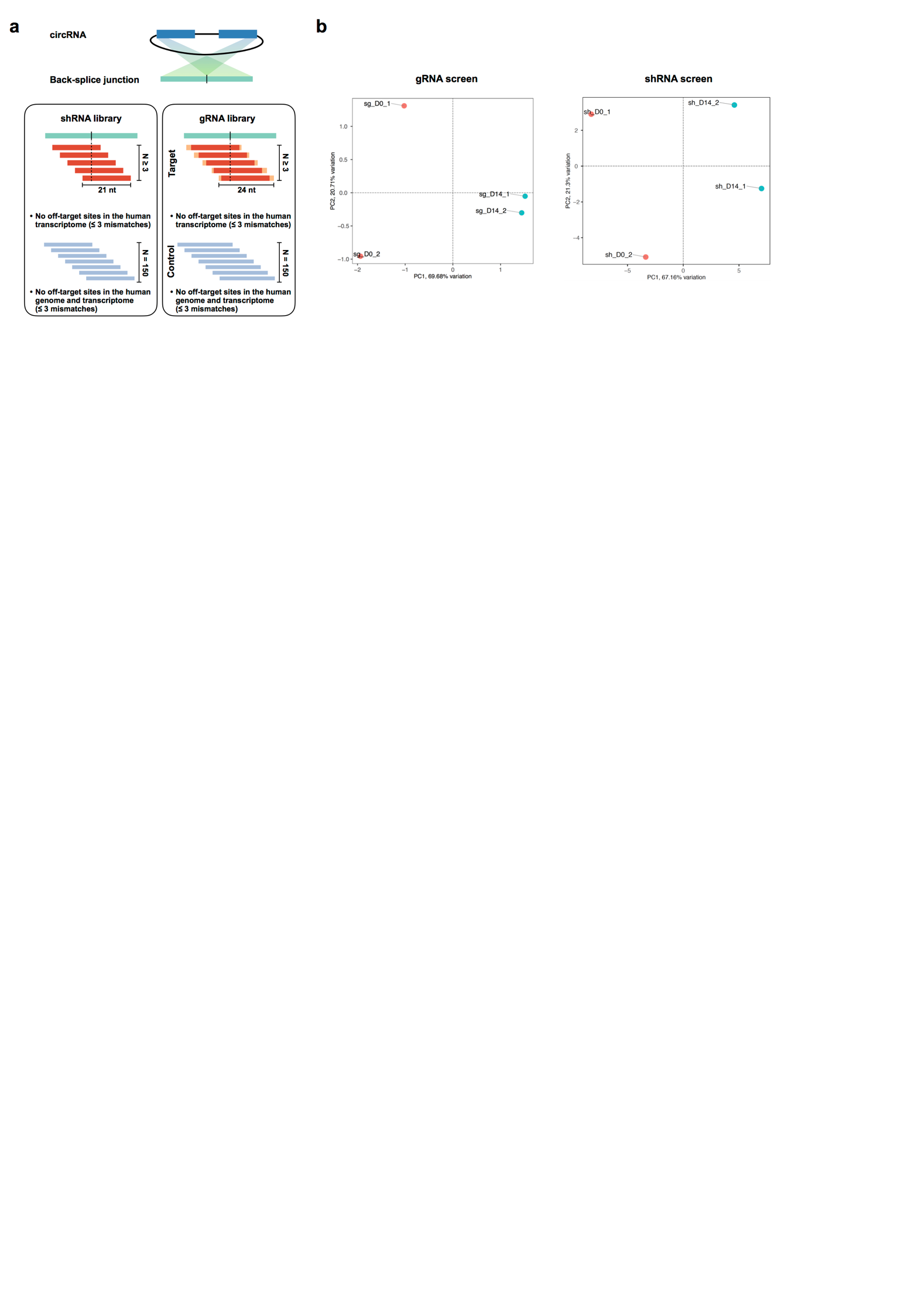

**Supplementary Fig. 4 | Design of Cas13d and shRNA libraries. (a)** Schematic drawing of shRNA and gRNA libraries (see details in Methods). Position-matched shRNAs and gRNAs were designed to target the BSJ sites of 134 circRNAs. shRNAs and gRNAs targeting mRNAs of 10 essential genes serve as positive controls. 150 non-human genomic sequence targeting shRNAs and gRNAs are included as negative controls. (**b**) Principal component analysis of gRNA (left) or shRNA (right) levels across the four generated sequencing libraries. PC, principal component.

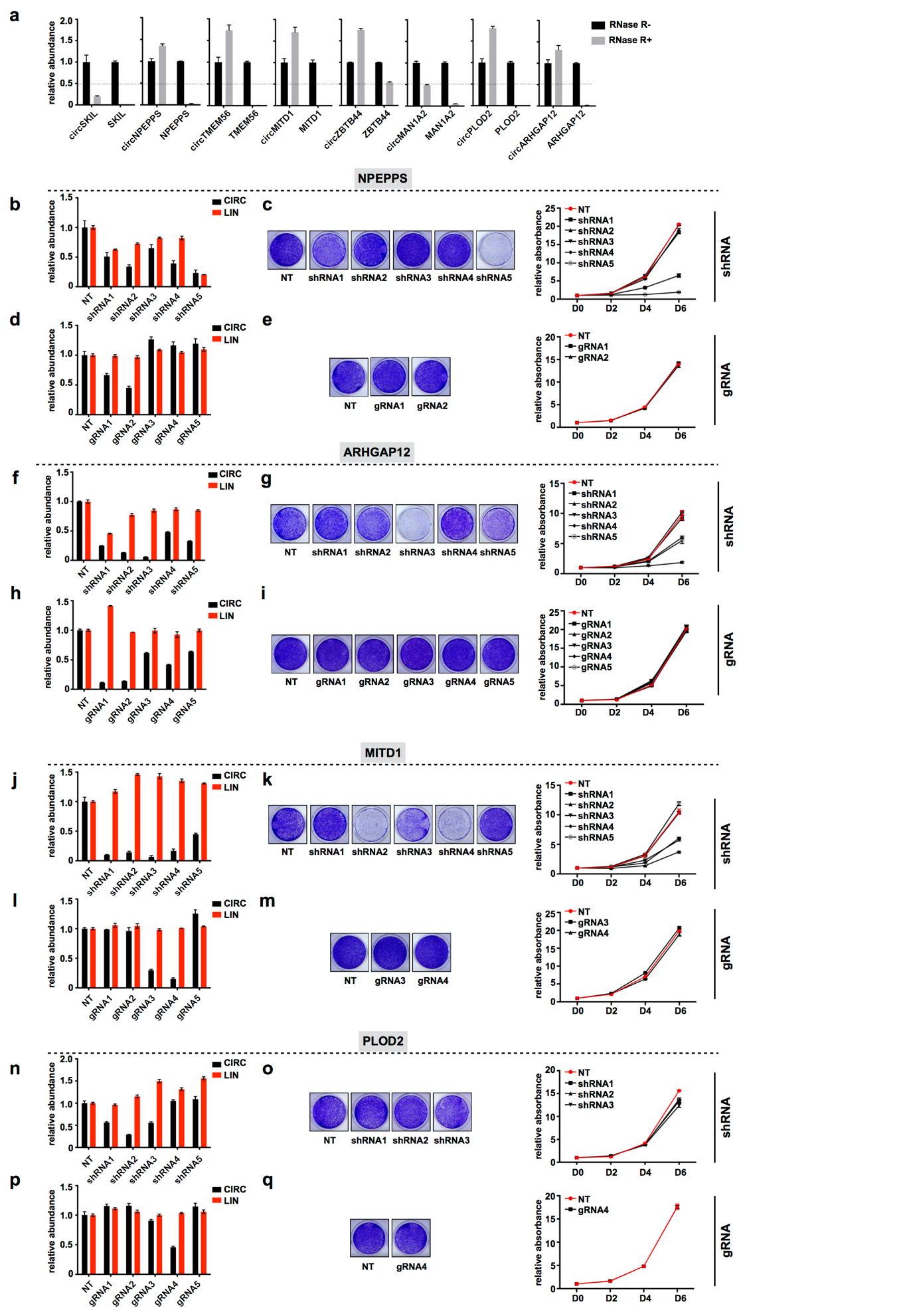

**Supplementary Fig. 5 | High false positive rate of shRNA functional screen for circRNAs. (a)** RNase R validation of 8 selected circRNAs. circRNAs together with the corresponding linear mRNAs were amplified by qRT-PCR from cDNA prepared from RNA non-treated or treated with RNase R. **(b, d)** Relative expression levels of circNPEPPS and its parental mRNA upon knock-down of circNPEPPS by shRNAs (b) and gRNAs (d) in Huh7 cells. **(c, e)** Proliferation rates of control and shRNA-mediated (c) and Cas13d-mediated (e) circNPEPPS-silenced Huh7 cells. The number of cells was detected upon staining with crystal violet, and representative pictures are shown on the left, while the proliferation curves are shown on the right. **(f, h, j, l, n, p)** same as in (b, d) for circARHGAP12, circMITD1 and circPOLD2. **(g, i, k, m, o, q)** same as in (c, e) for circARHGAP12, circMITD1 and circPOLD2. The data shown are from one of two biological replicates with similar results, and error bars indicating the mean ± s.d. of three technical replicates. **p* < 0.05, ***p* < 0.01, ****p* < 0.001 (unpaired student’s *t* test). ns, not significant.
